# Dynamic causal modeling for calcium imaging data reveals differential effective connectivity for sensory processing in a barrel cortical column

**DOI:** 10.1101/509653

**Authors:** Kyesam Jung, Jiyoung Kang, Seungsoo Chung, Hae-Jeong Park

## Abstract

Multi-photon calcium imaging (CaI) is an important tool to assess activity among neural populations within a column in the sensory cortex. However, the complex asymmetrical interactions among neural populations, termed effective connectivity, cannot be directly assessed by measuring the activity of each neuron using CaI but calls for computational modeling. To estimate effective connectivity among neural populations, we proposed a dynamic causal model (DCM) for CaI by combining a convolution-based dynamic neural state model and a dynamic calcium ion concentration model for CaI signals. After conducting a simulation study to evaluate DCM for CaI, we applied it to an experimental CaI data measured at the layer 2/3 of a barrel cortical column that differentially responds to hit and error whisking trails in mice. We first identified neural populations and constructed computational models with intrinsic connectivity of neural populations within the layer 2/3 of the barrel cortex and extrinsic connectivity with latent external modes. Bayesian model inversion and comparison shows that a top-down model with latent inhibitory and excitatory external modes explains the observed CaI signals during hit and error trials better than any other model, with a single external mode or without any latent modes. The best model also showed differential intrinsic and extrinsic effective connectivity between hit and error trials (corresponding to the bottom-up and top-down processes) in the functional hierarchical architecture. Both simulation and experimental results suggest the usefulness of DCM for CaI in terms of exploration of the hierarchical interactions among neural populations observed in CaI.

## 1. Introduction

In terms of exploration of the neural interactions within a cortical column, multi-photon calcium imaging (CaI) is an important tool to assess activity of neural population. Not only providing information regarding distributions of neurons (in superficial layers), CaI also allows exploration of the functional tuning of each neuron while perceiving a stimulus (Ohki et al., 2005) or conducting a task (Guo et al., 2014; Peron et al., 2015), with a relatively high temporal resolution (Grewe et al., 2010; Helassa et al., 2016). Using CaI, we are also able to investigate complex interactions among multitudes of neurons within a cortical column in detail, which otherwise is not generally possible.

A cortical column, as a functional unit of sensory perception, is understood to process bottom-up signals by integrating them with top-down information via the populations of excitatory neurons and inhibitory interneurons distributed across layers of the cortical column. The functional architecture of these processes is reflected in recent canonical microcircuitry models that include interactions within three or four neural subpopulations in the cortical column, in association with externals (Bastos et al., 2012; Douglas and Martin, 1991; Haeusler and Maass, 2006). These populations are considered to interact with others asymmetrically through feed-forward and feed-backward connections (Chaudhuri et al., 2015; Friston, 2018; Mesulam, 1998; Park and Friston, 2013), which is supported by neuroanatomical studies (Markov et al., 2014). Since the interactions among neural populations within a neural column (i.e., intrinsic connectivity) and those with neural populations outside the column (extrinsic) are generally reciprocal and asymmetrical, they can be better described by effective connectivity than by functional synchrony (Friston, 1994). However, the effective connectivity among neural populations, occurring at the cortical column in the sensory cortex, cannot be directly assessed by measuring the activity of each neuron using CaI but calls for computational modeling.

In the exploration of the neural interactions using computational modeling, CaI has innate limitations due to the transmission limit of light and the limitation in the spatial coverage for a given spatial resolution; CaI can primarily measure indirect neural activity in the superficial layers but not in the deep layers in vivo. Furthermore, CaI cannot measure activity across multiple columns at a single-neuron level (i.e., trade-off between coverage and spatial resolution). Since the cortical column itself does not act alone but integrates signals received from outside the column, we cannot fully understand the activity and interactions among neurons within the column through CaI measurements at a shallow portion (usually the layer 2/3) of a single column. The lack of measurements at the deeper layers and at regions outside the column demands estimation of hitherto unobserved data (such as hidden states or latent variables) under constraints of coevolving hidden-state, and connectivity estimation through observed data.

The purpose of the current study was to propose a method to explore effective connectivity among neural populations or individual neurons within a barrel cortical column using CaI, and to test the need for external modulation in a column of the barrel cortex during a task. For modeling columnar connectivity using CaI, we modeled the neural encoding at the mesoscopic level by clustering neurons into several groups according to the waveforms of their CaI signals. This approach is based on the *population coding* hypothesis, which states that the brain encodes sensory information in the neural population (Pasupathy and Connor, 2002; Pouget et al., 2000). It begins with estimation of the number of neural populations from a multitude of neurons. To estimate effective connectivity among neural populations, we proposed a type of dynamic causal model (DCM) (Friston et al., 2007; Friston et al., 2003) for CaI by combining a convolution-based neural state model modified from Moran et al. (2013) and a realistic CaI observation model (Rahmati et al., 2016). The proposed approach differs from the model of Rosch et al. (2018), where a fixed kernel function is used for CaI observation.

We applied the proposed modeling scheme in the exploration of integrative processing within a column in the barrel cortex using a public dataset (https://crcns.org/data-sets/ssc/ssc-2/about-ssc-2). The dataset contains CaI data of a column of the mouse barrel cortex during a pole localization task, responding to changes in the curvature and angle of a whisker in the localization step, which involved licking a reward port to receive water drops (Peron et al., 2015). In the original paper, Peron et al. (2015) reported that the proportions of representative neurons for touch and whisking, and their characters are stable during learning. Moreover, the touch and whisking behaviors showed different neural activity maps in a barrel cortical column. The authors concluded that the neurons in the layer 2/3 of the barrel cortex are organized to encode sensory inputs from whiskers. However, they did not fully explore how neural populations in the layer 2/3 of the barrel cortex respond differently to two different types of behaviors, i.e., those during hit and error trials. Thirsty mice successfully lick a reward port to receive water drops in the hit trials but failed to lick a correct port even though they touched the target pole in the error trials. In the error trials, the sensory input did not lead to rewards.

What is the difference in the effective connectivity among neural populations when the sensory cortex processes perception to successfully receive rewards or fail? The answer to this question may lie in the top-down and bottom-up interactions among neural populations in a cortical column. Specifically, the difference in success or failure would be reflected in the intrinsic effective connectivity among neural components in a column with differential extrinsic connectivity with externals. To test this hypothesis, we constructed and evaluated computational models with intrinsic connectivity of neural populations within the layer 2/3 of the barrel cortex and extrinsic connectivity with latent external sources (corresponding to the bottom-up and top-down processes).

The current paper is composed of four parts: 1) processing of CaI signals and extraction of neural populations; 2) introduction of DCM for CaI and explaining DCM models of neural populations for hit and error trials; 3) simulation and validation of DCM for CaI; and 4) evaluation of models, followed by the discussion on the use of computational models for explaining hits and errors in the perspective of extrinsic signaling within the layer 2/3 of a barrel cortical column.

## 2. Methods

### 2.1. Experimental Data

We analyzed functional CaI data of neuronal responses in the barrel cortex evoked by a single whisking cycle (Guo et al., 2014; Peron et al., 2015), which is available from a public database (https://crcns.org/data-sets/ssc/ssc-2). Briefly explaining the data, six 6–8-week old emx1-Cre X LSL-H2B-mCherry mice, which had red nuclei in excitatory neurons and glial cells (Gorski et al., 2002), were used. They were infected with AAV2/1 syn-GCaMP6s (Chen et al., 2013) to indicate the concentration of cytosolic calcium ions in neurons; CaI data from three 600 × 600 μm^2^ (512 × 512 pixels) imaging planes, separated by 15 μm in depth, were acquired at 7 Hz while each pre-trained mouse performed a task. In the task, a target pole appears within the ambit of whisking of a single whisker. Posterior and anterior locations of the target pole indicate the right and left lick-ports which supply water drops as rewards. The task was to select a lick-port that will supply water drops in a given trial. The hit trial indicates that the thirsty mice successfully chose a lick-port which supplies water drops. During the error trials, the mice failed to choose the correct lick-port. The task and data have been described in detail by Guo et al. (2014) and Peron et al. (2015), and preprocessing of CaI signals has been described by Huber et al. (2012). To examine trial type-dependent effective connectivity among the neural populations in a barrel column, we divided the trials into hit and error types and conducted computational modeling for these two performances.

The procedure for signal processing for computational modeling is presented in Figure 1 and is explained in the following sections.

**Figure 1.**
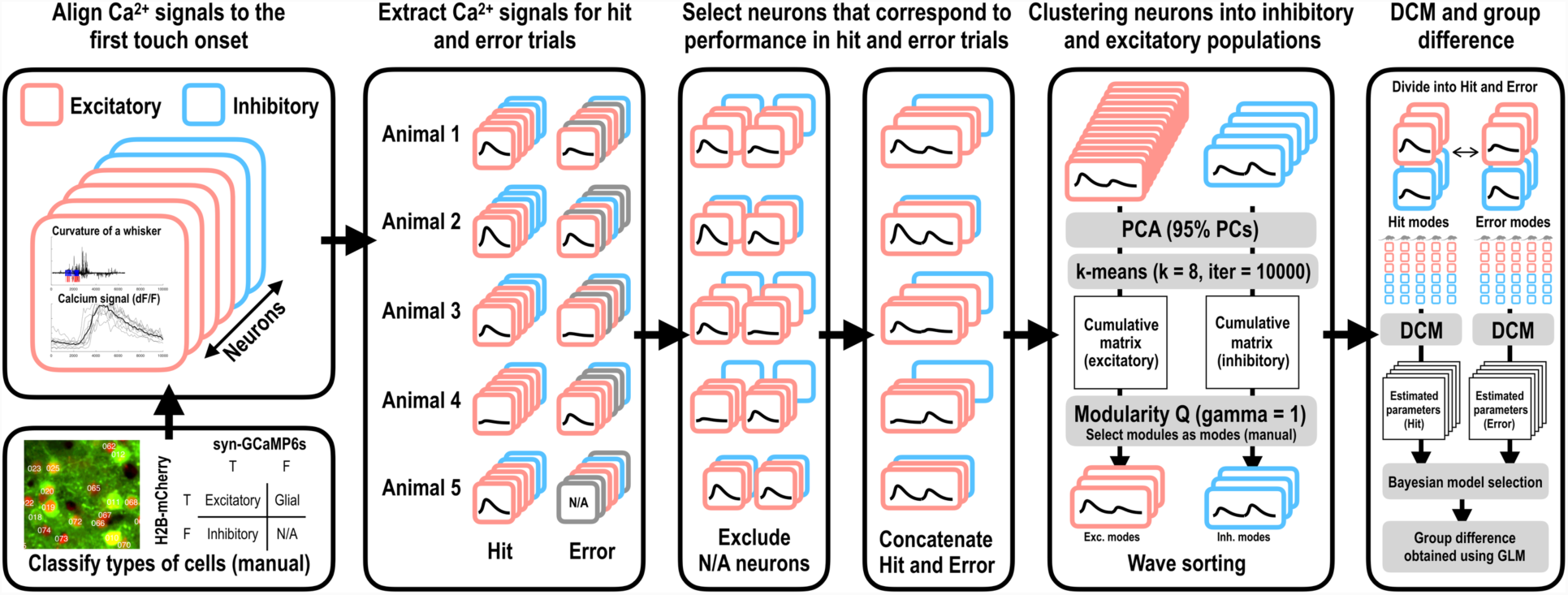
Procedure for signal processing and dynamic causal modeling for calcium imaging. It consists of waveform sorting, clustering signals according to hit and error trials and according to inhibitory and excitatory neural populations, and dynamic causal modeling to estimate the difference in effective connectivity for hit and error trials at the group level. N/A: not available, PCA: principal component analysis, Exc.: excitatory, Inh.: inhibitory, DCM: dynamic causal modeling, and GLM: general linear model

### 2.2. Classification of Inhibitory and Excitatory Cells

We selected responsive neurons based on the empirical criterion that the duration of high-signal changes (*ΔF/F*) (higher than 1) should exceed at least 5% of the total data points (see Figure 2). We then classified inhibitory and excitatory neurons based on red fluorescence channel images (mCherry; 675/70 emission filter, Chroma). A cell segment (provided in the dataset) was identified as excitatory if the intensity of pixels inside the target cell segment (H2B-mCherry) was brighter than its surroundings (neuropil), and was otherwise considered inhibitory. Using this criterion, 595 excitatory neurons and 151 inhibitory neurons were manually identified from all the five mice.

**Figure 2.**
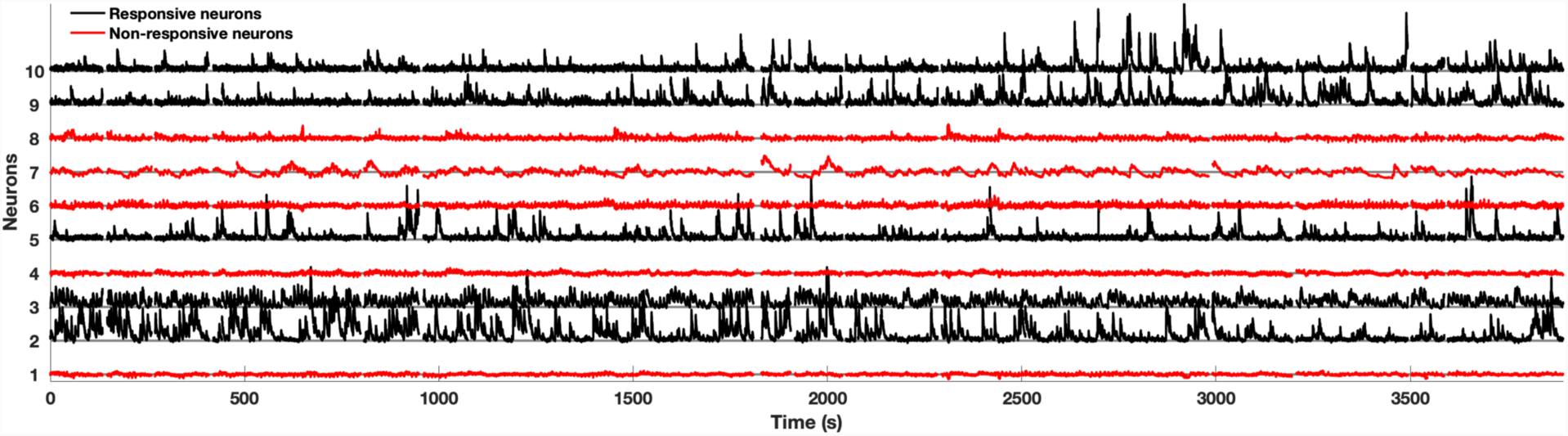
Calcium signals (*ΔF/F*) across a whole session. Black lines represent signals for selected neurons that show responses during whole trials. Red lines represent signals for neurons that are not active for the task.

### 2.3. Identification of Neural Populations

We aligned the calcium signals for each neuron to the first touch onset time for every trial, and divided the aligned signals into hit-right (hit the pole while whisking toward the right) and error-right groups. Since the mice employed a whisking strategy that maximizes touches for hit-right trials (corresponding to the posterior pole position) and minimizes touches for hit-left trials (Guo et al., 2014), we analyzed hit-and error-right trials for computational modeling as sensory input due to touches. After concatenating signals of hit and error trials for each neuron, we applied a moving average filter (14 Hz, cf. the sampling rate of the calcium signal was 7 Hz). In order to identify neural populations working synchronously (often termed modes), *principal component analysis* (PCA) was applied to the filtered signals of excitatory and inhibitory neurons (dimension: N_e/i_ neurons × T time series) to reduce temporal dimensions. The number of neural populations was determined using the following approach: 95% explainable principal component (PC) weights (for concatenated hit and error trials, dimension: P) for N_e/i_ neurons obtained by PCA (N_e/i_ × P) were used for *k-means* clustering with empirical K = 8 as the initial number of clusters for excitatory and inhibitory groups. To find a reliable number of neural populations, we repeated k-means clustering 10000 times for the PC weights. We then evaluated the frequency of identifying each pair of neurons as belonging to the same class according to every pair of k-means clustering (among 10000 executions). This composed a frequency adjacency matrix. For the frequency adjacency matrix, the *modularity optimization* algorithm (Newman and Girvan, 2004) in the Brain Connectivity Toolbox (Rubinov and Sporns, 2010) (Figures 3c and 3d) identified 5 and 4 modules for excitatory and inhibitory neurons while maximizing the modularity index (Q).

**Figure 3.**
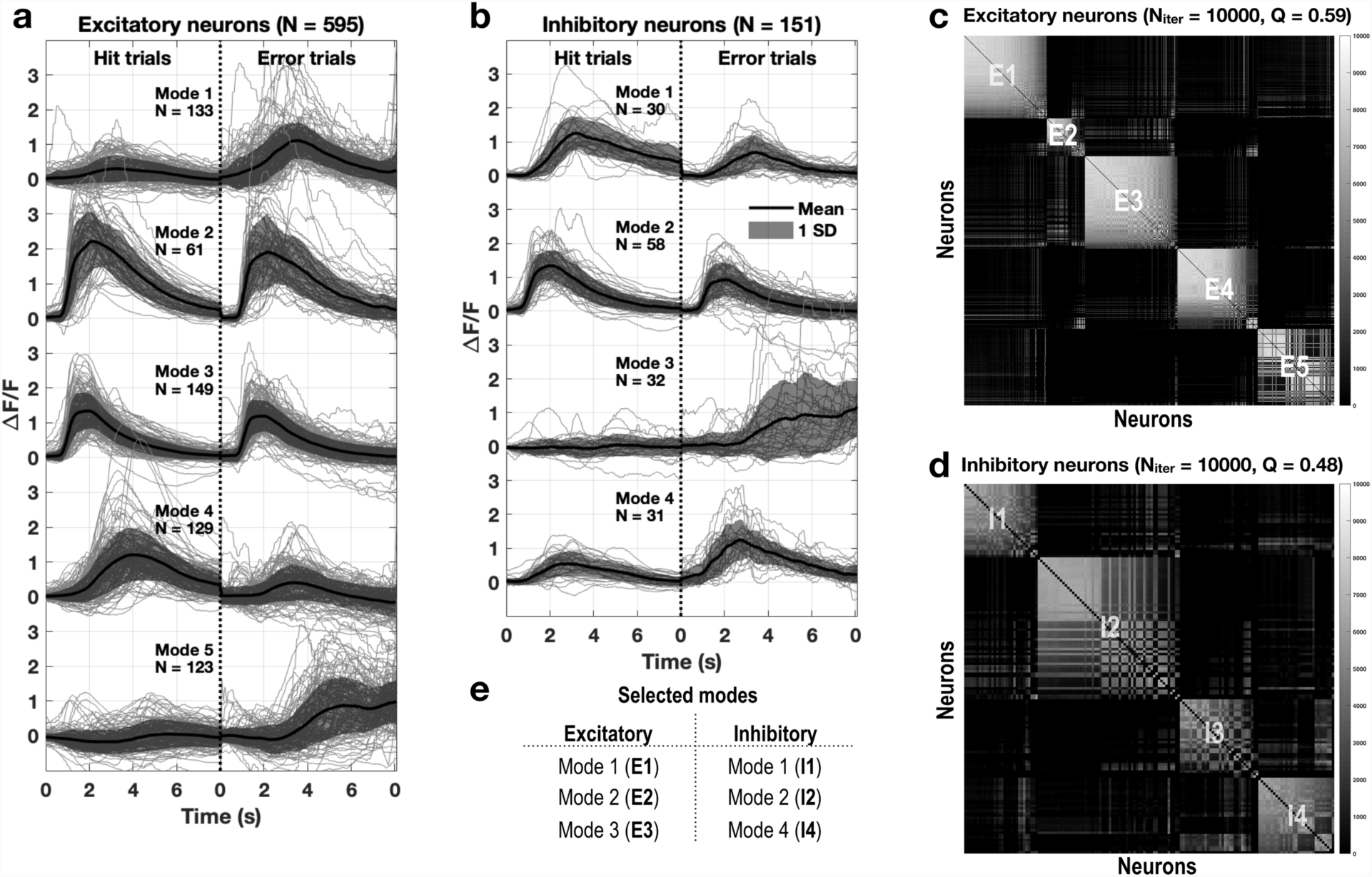
Wave-sorting and identification of neural populations. (a) Division of 595 excitatory neurons into five clusters by wave-sorting. Modes 1–3 were included for analysis. Although modes 4 and 5 showed response to the stimuli, they were rejected due to the delays of approximately 5 seconds after stimuli onset. (b) Division of 151 inhibitory neurons into four modes. Modes 1, 2, and 4 were included for analysis. Mode 3 was rejected due to its delayed response. (c and d) Excitatory and inhibitory frequency adjacency matrices show the frequency of the same cluster decided by pairs of the repeated k-means clustering process. The sequence of neurons was reordered based on the modularity optimization, which results in E1–E5 and I1–I4.

### 2.4. Dynamic Causal Modeling: Neural State Model

In order to construct computational models of interactions among the neural populations observed in CaI, we followed a DCM framework (Friston et al., 2007) implemented in SPM12 toolbox (https://www.fil.ion.ucl.ac.uk/spm/). We used a convolution-based model (Jansen and Rit, 1995; Moran et al., 2013) to describe the state dynamics of each neural population. In the convolution-based model for the neural state, a sigmoid activation function *σ*(*v*) converts a membrane potential to a firing rate. Then, the firing rate influences a post-synaptic potential through a synaptic kernel. Figure 4 illustrates the dynamics of the states of three neural populations (or modes) throughout a convolution-based model. The convolution-based model can be expressed as the following ordinary differential equation of cross-membrane current at a node (here, we explain it at a neuronal level, but it can similarly be used to describe the dynamics of neural populations, i.e., modes):

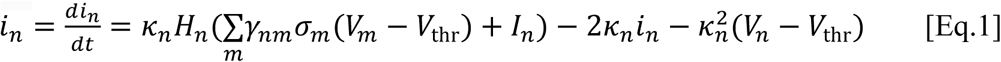

**Figure 4.**
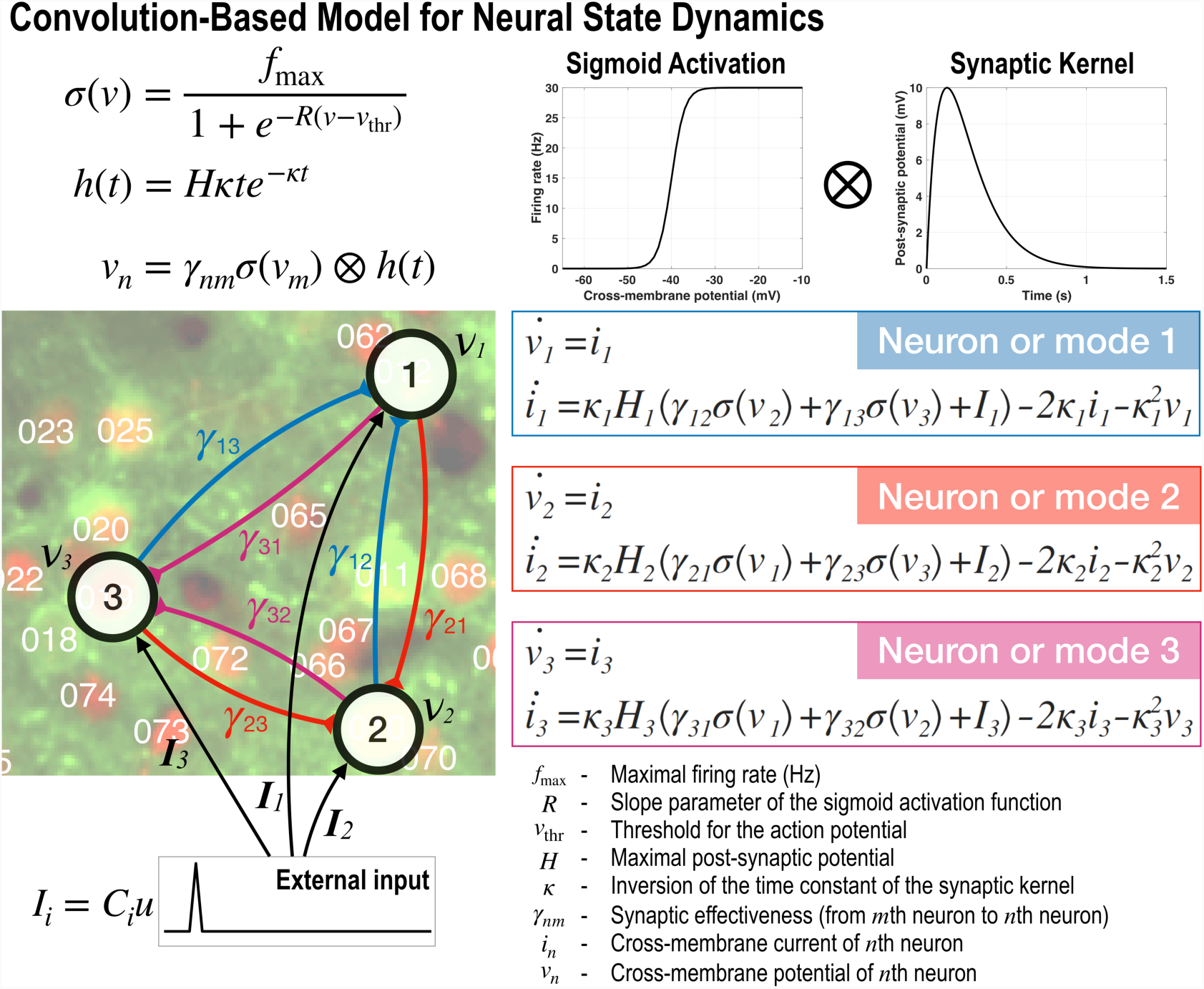
Computational model for calcium signal generation at a neural circuit composed of three exemplary nodes. The equations describe the convolution-based model for the neural state dynamics. The sigmoid function *σ*(*v*) stands for the relationship between the membrane potentials and the firing rates. The synaptic kernel *h*(*t*) transforms the firing rate into the post-synaptic potentials. The membrane potential *v*_*n*_ of a post-synaptic neuron is derived by convolution of the sigmoid function and synaptic kernel. The differential equations for the three nodes are derived by the convolution between the sigmoid activation function and the synaptic kernel as inputs. Description of the parameters used in the equations is presented at the bottom right side of the illustration.

Here, *i*_*n*_ is the cross-membrane current of a post-synaptic neuron; *V*_*m*_ and *V*_*n*_ are the cross-membrane potentials of the pre-synaptic (*m*) and post-synaptic neurons (*n*), respectively. *V*_thr_ is the threshold for action potential, *I*_*n*;_ is an external input on a post-synaptic neuron, and *σ* is a sigmoid activation function to convert the membrane potential to a firing rate. *κ* represents the inversion of the time constant of the synaptic kernel, and *H* is the maximal post-synaptic potential. *γ*_*nm*_ is the effective connectivity from all pre-synaptic neurons (all neurons denoted as *m* that have connections to the neuron *n*) to the neuron *n*. The cross-membrane potential for each neuron is updated by the following equation:

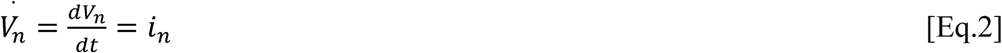

The assumption in Eq.2 is that the cross-membrane current equals the change in the cross-membrane potential if we consider the membrane capacitance to be a constant value (Hodgkin and Huxley, 1952).

We differentiated excitatory and inhibitory neurons in the computational model. The excitatory and inhibitory neurons induced depolarization and hyperpolarization of membrane potentials of the target neurons, respectively. As mentioned above, these dynamic equations can similarly be applied to the dynamics of neural populations (or modes), considering a neural population (a mode) to be equivalent to a neuron, as done in the current study.

### 2.5. Dynamic Causal Modeling: CaI Observation Model

The observation model for CaI was composed of a differential equation regarding the concentration of calcium ions ([Ca^2+^]) and a transformation from the calcium ion concentration to the calcium signal (Rahmati et al., 2016). The calcium dynamics (see Figure 5) can be described by the following equation:

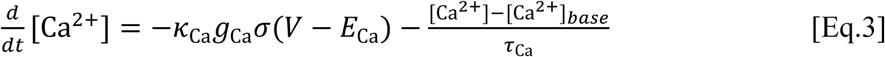

**Figure 5.**
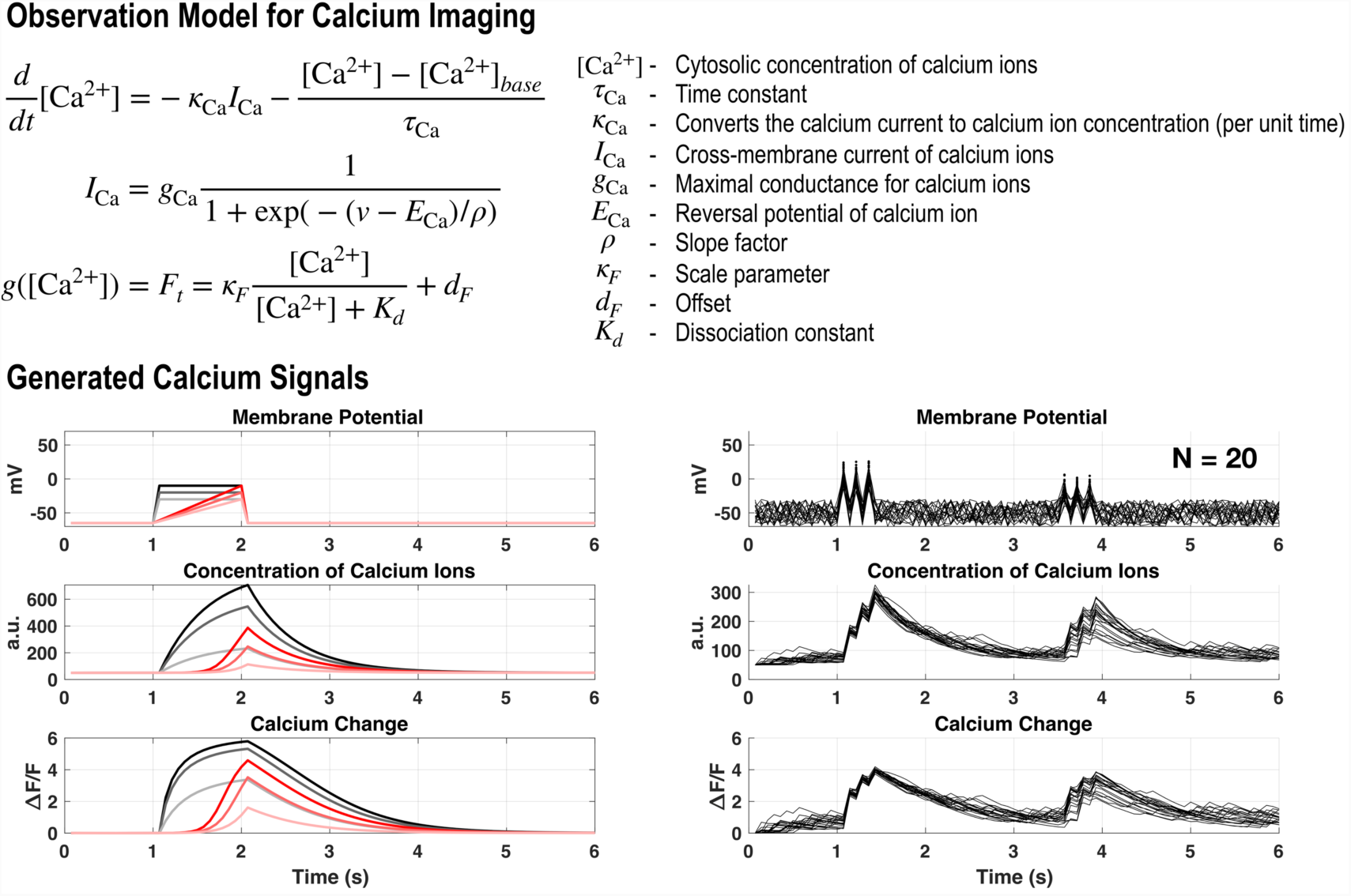
An observation model for calcium signals and its simulation examples. The top shows the equations for describing calcium dynamics, with descriptions of the parameters used. The bottom shows the results of two simulations using the observation model for calcium imaging. The bottom left panel displays the changes in the concentration of calcium ions and corresponding signals (*ΔF/F*) due to simple changes in the membrane potential while the right panel presents those for more complex changes in the membrane potential, with a background noise (N = 20).

Here, the parameter *κ*_Ca_ is the conversion ratio from a calcium ion current to its concentration; *g*_Ca_ represents the maximal conductance of calcium ions, *E*_Ca_ is the reversal potential of calcium ion, and *τ*_Ca_ is a time constant. The transformation from a calcium ion concentration to a calcium signal is described by the following equation:

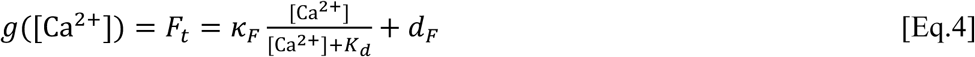

where *κ*_*F*_, *K*_*d*_, and *d*_*F*_ are scale, dissociation, and offset parameters, respectively. The parameters in Eq.3 and Eq.4 that describe calcium dynamics were obtained from previous studies (Rahmati et al., 2016; Yaksi and Friedrich, 2006). Figure 5 presents a simulation result for the observation model for CaI used in the current study. When different amplitudes and shapes of stimulations were applied to the resting membrane potential of a neuron, changes in the concentration of calcium ions and CaI signal are generated according to differential inputs.

### 2.6. Computational Models for a Barrel Column

The sensory processing in the layer 2/3 is known to integrate bottom-up afferent inputs from the layer 4 (L4, excitable mode), which is evoked mainly by thalamo-cortical signals (Petersen and Sakmann, 2001), and the top-down inputs (excitable and inhibitable) from cortical regions outside the cortical column. We hypothesized that the difference in the hit and error trials is associated with differential processing of intrinsic and extrinsic effective connectivity within the barrel cortical column.

To explain the mechanism of hit and error trials with respect to effective connectivity for top-down and bottom-up processing (Bastos et al., 2012; Douglas and Martin, 2004; Zagha et al., 2016), we constructed four computational models. All the models consisted of four parts: six neural populations in the layer 2/3 of a barrel cortical column, the ventral posteromedial nucleus (VPM) of the thalamus, the L4, and excitatory and inhibitory external modes (E1–E3; and I1, I2, and I4). In these models, only neural populations in the layer 2/3 are observed. The L4 and two external modes (E and I) are latent states to be estimated. Note that the two latent external modes may not have identifiable neural sources but rather indicate equivalent modes to elicit excitatory and inhibitory influences on the intrinsic neural connectivity in the layer 2/3. The latent states were set to follow the same neural dynamics as that observed in the neural populations in the layer 2/3. We assigned unidirectional connections from the VPM to the L4 and from the L4 to the neural populations in the layer 2/3 as bottom-up processes.

We assumed that the inhibitory modes in the layer 2/3 do not influence the external modes. In order to increase the influence sensitivity of the layer 2/3 to external modes, the excitatory modes in the layer 2/3 were set to activate the external modes (E and I) exponentially (i.e., non-zero effect size on a latent mode by using *eγnm*). Without this exponential constraint, the activities of modes in layer 2/3 did not influence the external modes sufficiently in the current experiment.

Based on the relationship with external modes, we constructed and tested four plausible models, which are presented in Figure 6.

1. Isolated model: This model is composed of interactions among populations within a barrel cortical column (intrinsic connectivity only), without interaction with external modes (Figure 6a). This model does not contain top-down modulations.
2. Excitatory top-down model: This model includes top-down modulation by an excitatory mode in the external sources (Figure 6b).
3. Inhibitory top-down model: This model includes a bidirectional interaction with an inhibitory mode in the external sources (Figure 6c).
4. Full top-down model: The fully connected model includes connections with the excitatory and inhibitory modes in the external sources (Figure 6d).

**Figure 6.**
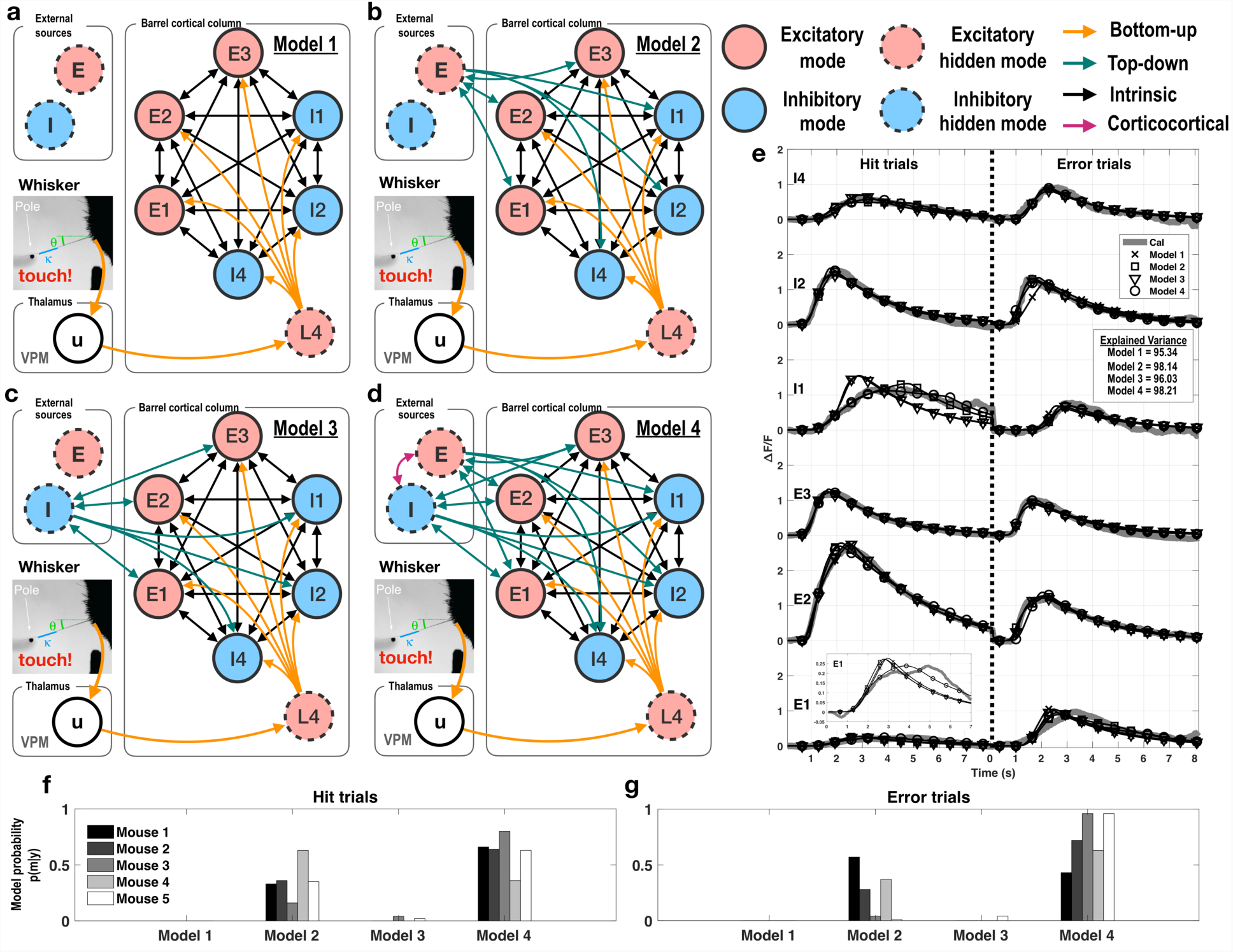
Four models for the barrel cortical column used in the current study. (a) Model 1 is a system of the interactions among six modes in the barrel cortex without interactions with external sources (isolated model). (b) Model 2 considers a latent excitatory external mode for excitatory top-down modulation. (c) Model 3 considers a latent inhibitory external mode for inhibitory top-down modulation. (d) Model 4 has two latent modes, as a fully connected model (excitatory and inhibitory top-down modulation). (e) As an example of model estimation results, observed calcium imaging signals and estimated signals are presented in thick gray lines and black-marker lines, respectively, for one mouse. (f and g) Model probabilities, p(m|y), for observed calcium imaging data, y, estimated for five mice, have been presented. VPM, ventral posteromedial nucleus

All these models generated CaI signals according to state transition and the CaI observation equations described above. The model probability for a given CaI datum was evaluated with respect to the log-evidence and conditional density for the parameters of each specific model. The best model was selected by comparing the evidences for four models using Bayesian comparison framework.

### 2.7. Bayesian Model Comparison and Parametric Empirical Bayesian Group Level Inference

To select the most compelling model among four models, we used a Bayesian model comparison in SPM12 (Stephan et al., 2009). This step compares the evidences after estimating the parameters of different models for the same data to identify the best model. After selecting the best model, we conducted group (between-subject)-level analysis for DCM of the selected model using the parametric empirical Bayesian (PEB) scheme in SPM12. This allowed us to identify between-subject effects based on specified design matrices at the second level, the details of which can be found in previous studies (Friston et al., 2015; Friston et al., 2016).

Briefly, the first-level (within-subject) effects are summarized in terms of posterior expectations and covariances, and are passed to the second-level (between-subject) to estimate posterior expectations and covariances of group means and between-subject effects. At the second level, the between-subject effects *β* on within-subject effects *θ* = {*A*, *C*, *θ*_*h*_} are encoded by a design matrix *X* (Eq.5 and Eq.6).

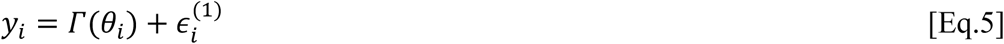

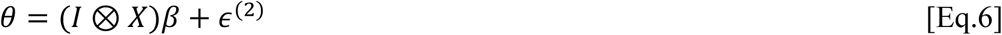

Here, *Γ* is a function that returns the predicted observations as a function of model parameters (corresponding to Eq.1 and Eq.2 for the neural state model, and Eq.3 and Eq.4 for the observation model). The hierarchical or PEB model used in the current study indicates that the parameters *θ*_*i*_of the *i*th subject are modeled as the sum of the group average and a random effect *∈*(2). In the current study, we used a design matrix *X* of seven regressors, which are group averages, a group (condition, hit vs. error) difference, and five hit-error pairs of five subjects (Figure 9). To reduce the number of parameters required to be estimated when using paired dependency (hit and error trials for each mouse), we simply did not consider within-subject dependency for group means of each trial types (Figure 8).

## 3. Results

### 3.1. Selection of Neural Populations via Wave-Sorting

The calcium signals of 595 excitatory and 151 inhibitory neurons in the layer 2/3 of the barrel cortex of five mice were sorted into multiple modes. When the modularity-optimizing process was applied (see Figure 1), the modularity Q of excitatory (eight modes) and inhibitory (five modes) clusters were 0.48 and 0.40, respectively (Figure 3), which indicate a strong community structure, and thus, a strong consensus across k-means clustering experiments (Newman and Girvan, 2004).

Among the 13 modes, those which had no evoked signals (less than 1 *ΔF/F*) or delayed peaks (over 4 seconds) were excluded from the current study. Finally, six modes (three excitatory and three inhibitory) were included for the computational modeling of the layer 2/3.

### 3.2. Model Parameters for Calcium Imaging

To make the models realistic, we used a set of fixed parameters for neural state and calcium signal dynamics based on previous studies (Moran et al., 2013; Rahmati et al., 2016; Yaksi and Friedrich, 2006). The parameters used in this study are listed in Table 1. The unfixed model parameters were estimated using the expectation-maximization process in DCM, with the parenthesized values in Table 1 as initial values. During the pre-stimulus period of each trial (zero CaI), the negative offset parameter (*d*_*F*_ in Eq. 4) for the fluorescence trace equals the signal change term of Eq. 4, *κ*_*F*_ [Ca^2+^]/([Ca^2+^ + *K*_*d*_]). Therefore, the value of the offset parameter was computed by using the scale parameter, baseline concentration, and dissociation constant.

**Table 1.** Parameters used in a computational model constructed for dynamic causal modeling of calcium signal

| Parameter | Variable | Type | Value* | Unit | Reference |
| --- | --- | --- | --- | --- | --- |
| Time constant of the kernel of PSP | $\kappa$ | Unfixed | (0.128) | ms | Yaksi et al., 2006 |
| Maximal PSP | $H$ | Fixed | 10 | mV | Moran et al., 2013 |
| Sigmoid activation function | $\sigma(V_{\text{pre}} - V_{\text{thr}})$ | Fixed | 0.67** | - | Moran et al., 2013 |
| Maximal firing rate | $f_{\text{max}}$ | Fixed | 30 | Hz | |
| Time constant of the kernel of calcium | $\tau_{\text{Ca}}$ | Unfixed | (0.8) | s | Rahmati et al., 2016 |
| Converting and scaling parameter | $\kappa_{\text{Ca}}$ | Unfixed | (15) | s<sup>-1</sup> | Yaksi et al., 2006 |
| Maximal conductance for calcium ions | $g_{Ca}$ | Fixed | 5 | mS/cm<sup>2</sup> | Rahmati et al., 2016 |
| Sigmoid calcium current function | $\sigma(V - E_{Ca})$ | Fixed | 0.2** | - | Rahmati et al., 2016 |
| Scale parameter for the fluorescence trace | $\kappa_F$ | Fixed | 10 | - | Yaksi et al., 2006 |
| Offset parameter for the fluorescence trace | $d_F$ | Fixed | -3.33 | - | (calibrated) |
| Baseline calcium ion concentration | $[Ca^{2+}]_{base}$ | Fixed | 100 | nM | Rahmati et al., 2016 |
| Dissociation coefficient | $K_d$ | Fixed | 200 | nM | Rahmati et al., 2016 |

\*Values in parentheses were used as initial values for parameter estimation.
\*\*Slope of the sigmoid function

### 3.3. A Simulation Study of the Computational Model

To test the validity of DCM for CaI data, we conducted a simulation study. We established a ground truth model with a pre-specified connectivity and parameters (Figure 7a), and generated membrane potentials and CaI signal changes in each neuron according to the dynamic neural state and observation equations. To accommodate variations within a group, we generated signals for each individual neuron using a set of parameters sampled under Gaussian distribution with the parameter values given in Figure 7a as mean and standard deviation, i.e. N(mean, variance). The box in Figure 7c shows an exemplary case of the simulation. For the generated CaI signals, we inverted three models specified using different connectivity configurations.

**Figure 7.**
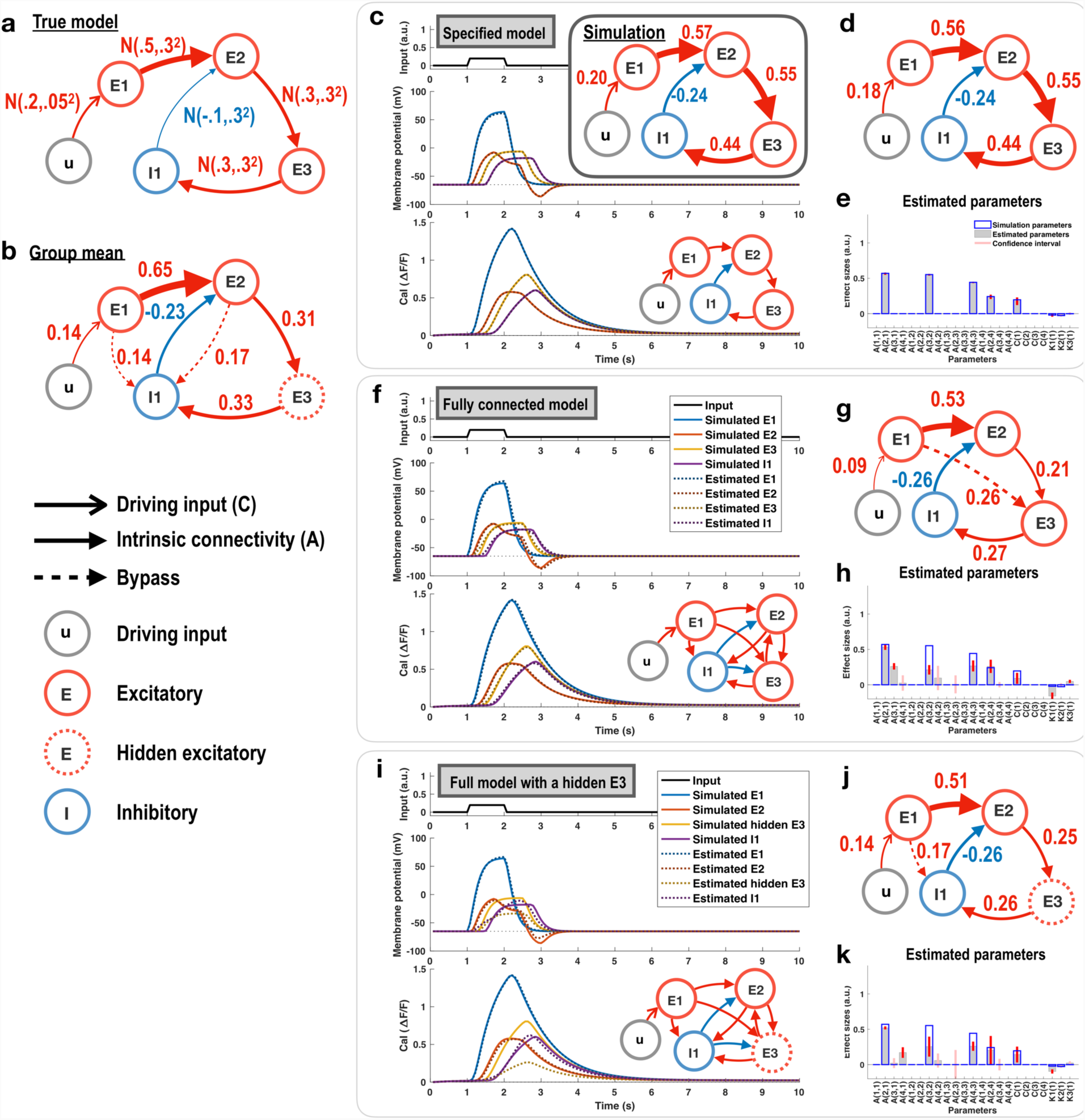
Simulation of the computational model for the calcium signals for 10 subjects. (a) A true model is shown. Effective connectivity estimated using a specified model; fully connected model; and fully connected model, observed at a neuron, with E3 as a hidden latent neuron, is shown. (b) Analysis performed using the parametric empirical Bayesian scheme displays a group-averaged effective connectivity. We have denoted the estimated connection which was absent in the true model (dashed arrows) as the “bypass” connection. (c) A set of parameters for each individual was sampled from a true model (a) under the Gaussian distribution. A driving input u activates E1 neuron (top). Simulated (solid lines) and estimated (dotted lines) membrane potentials (middle) and calcium signals (bottom) for each neuron are presented. The specified model shows reliable estimation of the parameters (d) compared with the values of simulation in (c). (d, g, and j) Effective connectivity estimated using a specified model; fully connected model; and fully connected model, observed at a neuron, with E3 as a hidden neuron, is shown. (e, h, and k) Expectation (blue bar) and 95% credible interval (red bar) for each parameter in each model, and the true parameter values are shown. (f) The parameter estimation performed using a fully connected model (with connections among E2, E3, and I1) is presented. (i) The parameter estimation performed using a fully connected model, but regarding E3 as a hidden neuron (not using the E3 activity data in the model inversion).

First, we specified a model with an identical intrinsic connectivity to the true model, and applied model inversion using DCM. It reliably estimated parameters of the specified model with high precision (Figure 7e). Membrane potentials and calcium signals for all neurons (E1, E2, E3, and I1) assessed using the estimated parameters are displayed in Figure 7c (dotted lines) with simulated signals (solid lines).

Second, since a prior knowledge of the intrinsic connectivity is not generally available, we assumed that all neurons (E2, E3, and I1) are fully connected with each other (a fully connected model), and estimated the parameters of this model using DCM. The group mean of all DCMs was estimated using PEB, and the parameters with a large 95% credible interval (that contains zero) were removed from the fully connected model (we term this a reduced model, Figure 7g). Figure 7f shows the generated CaI signals, and membrane potential resembled the true signals with high correspondence, but the effective connectivity (Figures 7g and 7h) was not identical to the ground-true effective connectivity. However, the difference can be considered a causal-chain effect (from E1 to E2, and from E2 to E3), such as a *bypass* (from E1 to E3). The effect size from E2 to E3 of the fully connected model was lower than that of the true model, but the bypass from E1 strengthens the target E3 activity. Consequently, E1 and E2 influence E3 as much as in the true model.

Third, we inverted a fully connected model with a hidden neuron (E3), simulating the lack of observed data for E3. The hidden neuron was controlled under the same computational model as the observed neurons. Most estimated CaI and membrane potential signals (except those for the hidden neuron) fit the signals generated using the true model. However, the estimated signals for the hidden neuron showed smaller amplitudes than the true signals but showed a time course similar to that of the true signal. The fully connected model with a hidden neuron also showed a bypass connection from E1 to I1.

### 3.4. Bayesian Model Comparison by Model Probability

To select the most reliable model among four models for the real CaI data set, a model evidence (probability) for a given datum for each subject was calculated using Bayesian model comparison. The results are displayed in Figures 6f and 6g. In both hit and error trials, the model with full excitatory and inhibitory connections (model 4) was found to be the best for describing CaI signals.

### 3.5. Effective Connectivity for Different Types of Trials

The parameters of the full excitatory-inhibitory model (model 4) for hit and error trials were estimated using DCM (see Figure 8), followed by application of the PEB scheme for the group analysis. We considered an effective connectivity as significant if it differed from zero with 95% posterior confidence.

**Figure 8.**
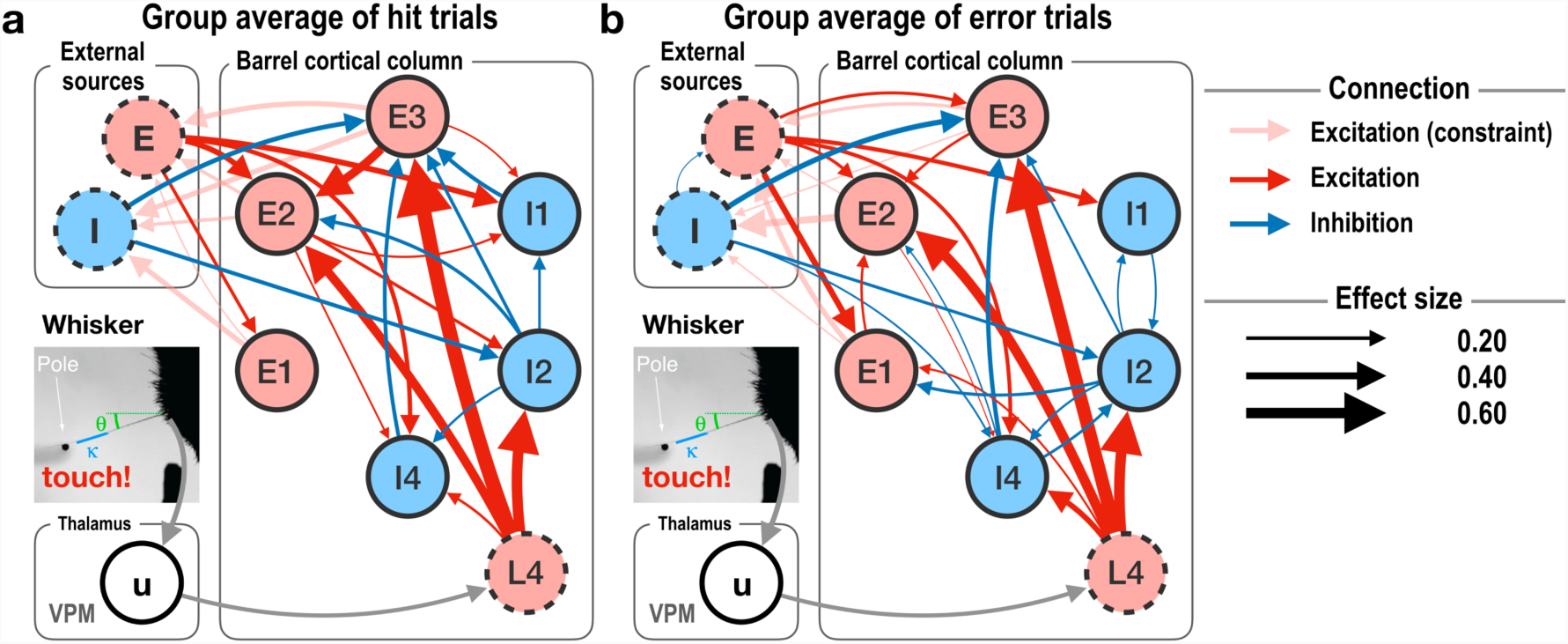
Results obtained using parametric empirical Bayesian scheme for assessing common group effective connectivity across mice for hit (a) and error (b) trials for the best model (model 4). (a and b) Group-averaged effective connectivity during hit and error trials are displayed. In both cases, the inhibitory modes exert a smaller influence on the other modes than the excitatory modes. The effect of excitatory signals from the excitatory modes in the barrel column cortex to the external sources were adjusted to ensure exponential sensitivity (constraint). VPM, ventral posteromedial nucleus

In the bottom-up process, the L4 mainly affects E2, E3, and I2 in both the trial-types (Figure 8). As mentioned in the simulation and validation sections, it is probable that a part of the L4-E3-E2 causal chain-connection may be split into bypass effect from the L4 to E2 (Figure 8a). Although this bypass may not be dissociable from the poly-synaptic connection from the L4 to E3, it shows the bottom-up direction of the information flow. In the top-down process, the excitatory latent external mode influences E1, E2, I1, and I4, while the inhibitory latent external mode suppresses E3 and I2.

The common and different effective neural connections between hit and error trials were evaluated using a group-level PEB (Figure 9). Figure 9a shows a common effective connectivity for hit and error trials. For both hit and error trials, there were bottom-up and top-down connections with the six modes in the layer 2/3. The bottom-up excitatory L4 affects mainly E2, E3, and I2. The excitatory external source modulates E1, E2, I1, and I4, and the inhibitory external source inhibits E3 and I2.

**Figure 9.**
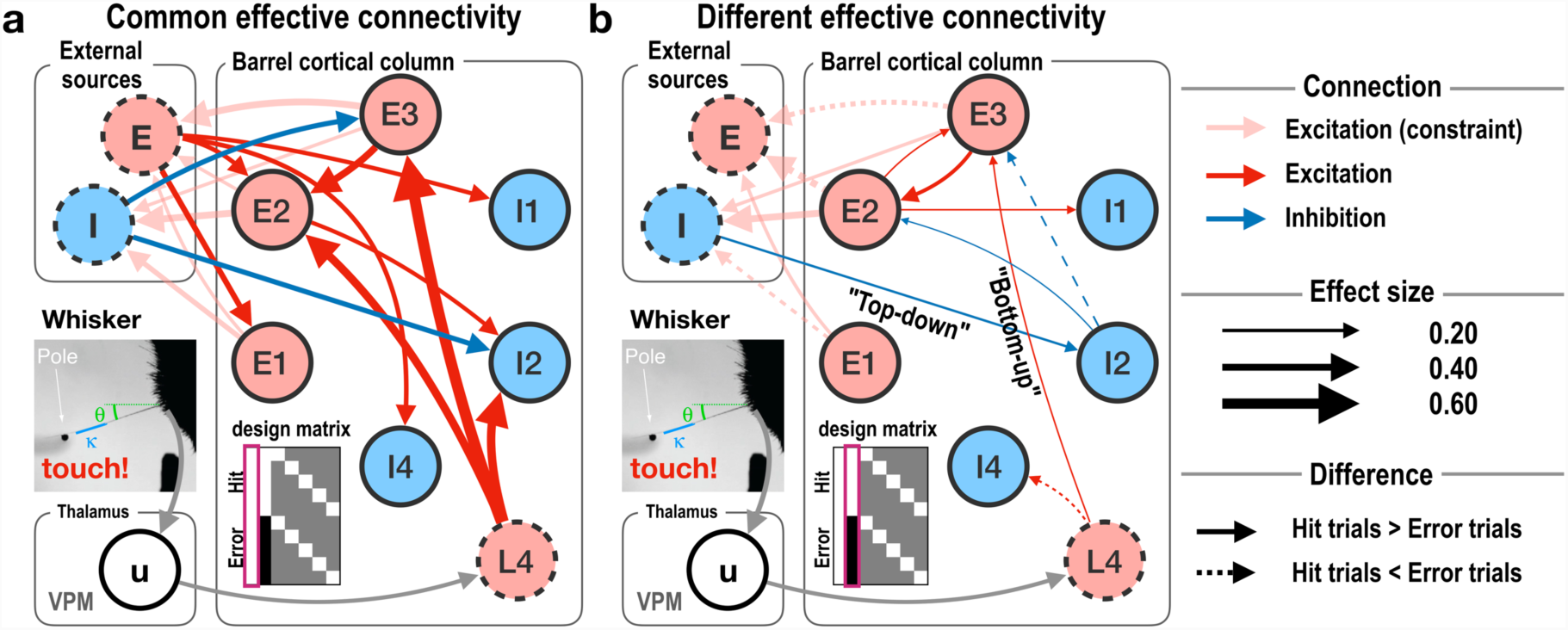
Results obtained using parametric empirical Bayesian scheme for assessing common and different effective connectivity between conditions. (a) The illustration shows the common effective connectivity between two types of trials (hit and error conditions). It corresponds to the first column (constant) of the design matrix. (b) The illustration displays difference in effective connectivity between error and hit trials (condition difference). It corresponds to the second column (hit minus error) of the design matrix X in Eq.6. The main difference exists in inhibition of the excitatory association cortex by the inhibitory external source. Consequently, the effectiveness of stimulation of the barrel cortex by the excitatory external source decreases during error trials. Red and blue colors indicate connectivity with the excitatory and inhibitory sources. Solid and dotted lines in (b) indicate increased and decreased connectivity, respectively, in the hit trials compared with the error trials. VPM, ventral posteromedial nucleus

The difference between hit and error trials in the effective connectivity is shown in Figure 9b. For the bottom-up process during error trials, the latent excitatory L4 influences I4 more, but E3 less, than during hit trials. This reduces the activity of E3 in error trials. Although I4 did not significantly affect E3, the group-averaged intrinsic connectivity of the two types of trials (Figure 8) showed that, in both types of trials, I4 inhibits E3. The bottom-up modulation due to the L4 suppresses E3 more during error trials than during hit trials. Consequently, E3 influences E2 less, and E2 influences E3 less, which may lead to reduced influence of the latent inhibitory external source. Therefore, the latent inhibitory external source inhibits I2 less during error trials than during hit trials. This may lead to more activity at I2, which may affect (inhibit) E3 to a greater extent. This recursive loop changed the intrinsic connectivity of the layer 2/3 in the barrel cortical column during error trials.

## 4. Discussion

In this study, we proposed a computational scheme for analyzing neural interactions reflected in the CaI under a DCM framework by combining neural state and calcium ion concentration dynamic models. Using this method, we evaluated intrinsic and extrinsic effective connectivity between neural populations in a barrel cortical column differentially responding to hit and error trials. Using the current computational model, we showed that the external signals outside the barrel cortical column play an important role in explaining hit-and error-related neural activity within the column.

### 4.1. Computational Modeling for Calcium Imaging

The change in cytosolic concentration of calcium ions is known to directly reflect neural activity (firing rate of action potential) (Berger et al., 2007). Since calcium signal indicates the discharge of post-synaptic action potentials, it is advantageous for evaluating causal relationships or effective connectivity among neurons or neural populations. To model effective connectivity using CaI data, we combined a neural state dynamic model and CaI observation model following the general framework of DCM. For the neural state dynamic model, we used a convolution-based model which comprises inter-modal connectivity with an activation function from membrane potential to firing rate and a post-synaptic potential kernel for each neural population (Moran et al., 2013). For modeling CaI signals, we adopted a model of calcium dynamics to formulate the cross-membrane calcium current gated by the cross-membrane potential (voltage-gated calcium channels) and a model for the kinetics of cytosolic concentration of calcium ions (Rahmati et al., 2016). This differs from the convolution-based approach used by Rosch et al. (2018), who applied an exponential decay kernel (with the parameters of GCaMP6f obtained from the study by Chen et al. (2013)) to local field potential reflected in the calcium signals for the analysis of seizure activity.

The computational model we proposed includes parameters for the kinetics of calcium signals that differ according to calcium indicators. For example, GCaMP6s used for CaI in the current study shows not only higher sensitivity (approximately sevenfold) but also has a nearly twice rise time (time-to-peak) and nearly threefold slower decay time than GCaMP6f used by Chen et al. (2013). Thus, we used a longer time constant for the kinetics of calcium signal than that of GCaMP6f. Some model parameters can be assigned as *fixed* constants or as variables to be estimated (Table 1), depending on the question of interest. The fixed parameters had been selected based on previous literature.

By conducting simulation experiments, we confirmed that the proposed DCM for CaI can reasonably estimate latent neural states and effective connectivity from CaI data. As part of the Bayesian inversion framework, we confirmed, by simulation, that more prior knowledge regarding the circuitry or model parameters leads to more reliable estimation results (Figures 7d and 7g). However, in practice, we often lack prior knowledge regarding the system configuration. In such cases, we can use Bayesian model comparison to select the best model to explain the observed data. We began our approach using a fully connected model without prior knowledge of the connectivity among nodes. The fully connected model can be compared with reduced models in terms of accuracy and complexity (the number of parameters), termed free energy (Friston et al., 2016).

We often need to estimate connectivity of the whole system using partially observed data. In the present study, from the simulation experiment, we also confirmed that DCM can invert neural states and connectivity with hidden unobserved nodes (Figure 7j). The fact that neurons or neural populations do not work in isolation but interact via a certain connection imposes constraints on each nodal activity. This compels us to unravel hidden (latent) nodal activity and connectivity while coevolving circuity estimation using observed data. In the current simulation, the optimization process in DCM determines effective connectivity (the parameters of a model), which was absent in the true model (Figures 7a and 7b). It is possible that the estimated parameters in the best model are not a unique solution for describing the observed data (Van Geit et al., 2016). A multitude of different configurations can generate the same functional activity, which is often termed degeneracy of neural systems (Edelman and Gally, 2001). In this case, model complexity matters in the selection of the best model, according to Occam’s razor theory (Sober, 1990). In our simulation, the best model, however, highly corresponds to the true model in terms of total effect on nodes, split into the poly-synaptic and bypass effective connectivity.

### 4.2. Population Activities and Their Effective Connectivity

Although for CaI data, DCM can be used to estimate interactions among individual neurons, we applied it to neural populations. Previous studies have shown that neurons in the brain encode and process information as populations rather than individually (Pasupathy and Connor, 2002; Pouget et al., 2000). This principle of population coding (or rate coding) provides a theoretical basis for developing a microcircuit or brain-like system in a practically accessible manner (Bastos et al., 2012; Mizrahi et al., 2018). In this type of neural population (or ensemble) coding approach, neurons in the population are assumed to share the same neural state and are not distinguishable from each other in terms of activity (Deco et al., 2008; Knight, 1972).

In the current study, the calcium signals evoked in a multitude of neurons in the layer 2/3 of a barrel cortical column were sorted and clustered into six neural populations. However, for modeling a barrel cortical column, we did not restrict the interactions among neural populations within the column. To determine whether to include external neural populations as latent modes (E and I), we constructed four models and tested them to identify the best model for explaining the activity observed in the six neural populations within the barrel column. The differentiation of neural populations was based on their neural activity pattern for external inputs. Thus, this approach does not differentiate the interconnection among neurons or consider neural diversity within the same population but abstracts them as a functionally unimodal node. This abstraction is advantageous as well as disadvantageous: advantageous in terms of simplifying the model and making computational modeling accessible for addressing population-level questions, disadvantageous in terms of explaining neurobiological properties and connectivity among neurons.

### 4.3. Latent States for Intermingled Modulation by Other Regions

Touching a whisker evokes receptive neural activity in a specific column of the barrel cortex. This specificity allows the functional mapping of a target whisker (such as the directional preference map) in the layer 2/3 of a barrel cortical column (Andermann and Moore, 2006). Despite columnar specificity of whiskers, the neural population in a column of the barrel cortex does not only process bottom-up signals but also integrates information from (or modulated by) neighboring columns (Petersen and Sakmann, 2001) and/or other brain areas (Manita et al., 2015; Zagha et al., 2013).

For explaining the observed data within the superficial layer of a barrel cortical column, we confirmed that the incorporation of interactions with other brain regions (not observed) is necessary. This was evidenced by the comparison of four different models, in terms of presence and absence of external sources, and in terms of E, I, and E-I (Figure 6). Probabilities of the four models suggested that the model with hidden excitatory and inhibitory modes as modulators for the layer 2/3 in the barrel cortical column (i.e., model 4) better explained the observed calcium signals than the other models (Figures 6f and 6g: bar plots); the excitatory model was second to the fully connected model (i.e., model 2).

According to a study on neuronal morphologies of the layer 2/3 and L4 within a barrel cortical column by Petersen and Sakmann (2001), the excitatory neurons of the layer 2/3 have dense dendrites at the same layer and axonal arborizations to lateral columns and a deep layer (layer 5). Excitatory neurons of the L4 also have dense dendrites at the same layer and axons forming an ascending column of input to the layer 2/3 pyramidal neurons. Based on these findings, we constructed the current models for the layer 2/3 in a barrel cortical column that include a hidden node corresponding to the L4. The hidden-node L4 is required to generate neural activity of spiny-stellate cells in the same layer of a barrel cortical column. Considering that sensation is delivered to a barrel cortical column via a thalamocortical pathway, we designed the bottom-up pathway, VPM-L4-layer 2/3 (Figure 6, yellow arrows).

In addition, we introduced latent excitatory and inhibitory modes for external sources outside the model of a barrel cortical column. It has long been debated whether external input to a cortical column is inhibitory or excitatory (Wang et al., 2013; Wang, 2002; Zagha et al., 2016). The current computational model suggests the need of both types of external sources, and preference toward excitatory regulation when we are to select only one type of external source from the neighboring or long-range projections. The two external sources represent mixtures of all possible external effects on the column of the barrel cortex. A cortical column in the barrel cortex receives/sends neural signals through long-range projections from the contralateral (Petreanu et al., 2007) and ipsilateral (Kinnischtzke et al., 2014; Manita et al., 2015; Petreanu et al., 2012) cortices. These long-range projections to the somatosensory cortex have mostly been studied with respect to the motor cortical feedback pathway. In the current model, external sources may include more than these long-range projections, including lateral connection with neighboring columns in the barrel or motor cortices. The details of the types of external sources and interconnections with them remain to be explored using more experimental data.

In the current study, we speculated that the external sources may primarily generate top-down modulatory signals that regulate the neural populations within a barrel cortical column. The top-down modulation during error trials was found to inhibit the intrinsic effective connectivity in the layer 2/3. Manita et al. (2015), for example, showed that the somatosensory cortex (S1) and secondary motor area (M2) form a top-down circuitry to perceive tactile stimulation. Interestingly, the study revealed that sensory stimulation induces neural activity sequentially from the S1 to the M2, followed by the M2 to the S1 in a reverse sequence. This sequential and reciprocal signal processing is similar to the processing of the best model in the current study. Moreover, the degradation of sensory perception by inhibition of the axons projecting from the M2 to the S1 is also similar to our results comparing effective connectivity between hit and error trials (Manita et al., 2015). However, this interpretation of external sources as playing a top-down role is a speculation that needs to be confirmed by more experimental data. Regardless of the role of external sources, the current results suggest that the modulation by external sources is necessary for describing the neural activities in the layer 2/3 of a barrel column during a localization task involving touches and whisking.

In summary, we (i) proposed a computational model for CaI signals using DCM, (ii) explored computational models to assess sensory perception through intrinsic and extrinsic connectivity, and (iii) examined the effective connectivity of the same neural circuit for different sensory perceptions (hit and error). We confirmed that a hierarchical architecture that has feed-forward and feed-backward connections is essential for formulating neural activity within a barrel cortical column for primary sensory perception. Both simulation and experimental results suggest the usefulness of DCM for utilizing CaI signals for the exploration of interactions among multitudes of neuronal activities observed in CaI or other imaging modalities.

## Acknowledgement

This research was supported by Brain Research Program through the National Research Foundation of Korea (NRF) funded by the Ministry of Science and ICT (NRF-2017M3C7A1049051). The authors thank the data distributor (https://crcns.org/data-sets/ssc/ssc-2) for sharing their experimental data.

